# Rapid volumetric reconstruction and tracking for Fourier light-field microscopy enables real-time calcium imaging in freely behaving *Hydra*

**DOI:** 10.64898/2026.09.17.752193

**Authors:** Raymond Adkins, Ryan Hausen, Jamie Noss, Gerard Lemson, Jonathon Howard

## Abstract

Fourier light field microscopy (FLFM) enables high-speed volumetric imaging by encoding multiple angular perspectives of a three-dimensional sample onto a single image. For this reason, FLFM is well-suited to sparse and rapidly evolving biological systems. To aid in the adoption of FLFM, we present OpenFLR, an open-source software framework for real-time volumetric reconstruction, three-dimensional particle tracking, and calcium image processing using FLFM. OpenFLR reconstruction is distributed as four interchangeable interfaces: a Python library, a command-line script, an interactive web application, and an ImageJ/micromanager plugin, so that the pipeline is accessible to both developers and bench biologists. Building on established Richardson-Lucy deconvolution, we use a hybrid experimental-computational PSF calibration strategy and a triangulation approach to tracking to extract particle positions in 3D directly from raw light field frames, bypassing reconstruction. We validate the complete pipeline on GCaMP6s recordings of freely behaving *Hydra vulgaris*, tracking sparse populations of neurons as they undergo large three-dimensional displacements.

## 1 Introduction

Organisms and tissues are three-dimensional, which poses a challenge for light microscopy. Examples include developing tissues, cell sorting in microfluidic channels, or freely behaving cells (Mertz 2019; Prevedel et al. 2014; Ahrens et al. 2013). In such systems, conventional two-dimensional widefield imaging is fundamentally limited: axial motion is difficult to distinguish from changes in fluorescent intensity and objects that overlap in projection but differ in depth cannot be differentiated. Accurate characterization of such systems requires imaging methods capable of resolving dynamics at high temporal resolution.

Scan-based volumetric imaging methods address the 3D imaging requirement but are temporally constrained by the time to sweep through the sample. Point-scanning methods offer high spatial resolution but are too slow to capture rapidly evolving systems. Light-sheet microscopy can provide faster volumetric rates and improved contrast but still requires either sequential plane illumination or complex multi-objective arrangements. In each case the volumetric frame rate is set by the number of planes that must be acquired, placing an upper limit on the speed of the dynamics that can be followed.

Alternate approaches become available when the fluorescence signal is sparsely distributed across the volume, as it is for calcium signals confined to cell bodies or for markers confined to the nucleus. For such systems, volumetric information can be captured within a single camera exposure, using single-shot microscopy (Levoy, 2006; Hua et al. 2021; Abrahamsson et al. 2013; Antipa et al. 2017). These methods capture all the information in the volume in each frame and are therefore much faster than z-scanning methods, at the cost of spatial resolution and computational reconstruction.

Fourier light-field microscopy (FLFM) is one such approach (Guo et al. 2019; Scrofani et al. 2017; Cong et al. 2017). By placing a lens array in the back focal plane of the microscope, FLFM simultaneously records multiple angular perspectives of the sample on a single detector. The relative displacements of features across the lenslets encode the axial position of each fluorescence source, enabling volumetric reconstruction without mechanical scanning. Because the acquisition rate is limited primarily by the camera exposure time, FLFM is particularly suited to sparse biological systems exhibiting rapid three-dimensional motion (Cong et al. 2017).

Despite these clear advantages, widespread adoption of FLFM has remained limited by the computational cost and accessibility of existing reconstruction software (Stefanoiu et al. 2019; Incardona et al. 2023).Volumetric reconstruction is typically performed using an algorithm similar to Richardson-Lucy (RL) deconvolution which is computationally expensive and available in unoptimized MATLAB or unpublished C++ pipelines (Stefanoiu et al. 2019; Incardona et al. 2023). Recent efforts to improve FLFM software have focused largely on accelerating the reconstruction step itself. Algorithmic approaches modify the deconvolution to converge in fewer iterations, for example through projection- estimation–accelerated Richardson-Lucy schemes (Wu et al., 2025), while multiview reformulations exploit the redundancy of the lenslet views to update the volume in ordered subsets, achieving millisecond-scale reconstruction for high-throughput light-field flow cytometry (Fu et al., 2026). Deep-learning methods can likewise reconstruct at video streaming rates but require retraining for each optical configuration and sample type and generalize poorly across noise levels and aberrations (Wang et al., 2021). These approaches share two properties: they remain focused on producing a full volumetric reconstruction and are typically released as standalone reconstruction code rather than as complete, user-facing analysis pipelines.

We present OpenFLR, an open-source software package spanning the complete Fourier light field microscopy workflow, from raw camera frames to biological measurements. First, we describe a hybrid experimental-computational PSF calibration strategy that combines the physical accuracy of bead-based measurements with the noise characteristics of an analytical model, enabling robust reconstruction under conditions where a single bead as the experimental PSF would be too noisy for iterative deconvolution. Second, we describe a GPU-accelerated, computationally optimized reconstruction implementation that achieves real-time volumetric reconstruction at acquisition frame rates on commodity hardware, without network training. Third, we describe a triangulation localization approach that extracts three-dimensional particle positions directly from raw light field frames via a batch FFT operation, bypassing reconstruction entirely for tracking applications. Finally, we package the reconstruction pipeline behind four interchangeable interfaces, a Python library, a command-line script, a web application, and an ImageJ/micromanager plugin, to lower the barrier to adoption for non-specialist users. Because our reconstruction is a clean, open, GPU-accelerated Richardson-Lucy implementation, alternate acceleration schemes can be incorporated directly as backends. More importantly, OpenFLR extends beyond reconstruction, providing an integrated, deployable workflow that carries data from calibration through tracking to per-cell activity in freely behaving, deforming animals. We demonstrate the utility of this framework through whole-animal calcium imaging of freely behaving *Hydra* expressing GCaMP6s and tdTomato in their neural cell line. On a Quadro P5000 GPU, we were able to fully reconstruct and track at an average of >2 fps, and on an H100 GPU, reconstruction jumps to >10 fps, which keeps pace with the rate required to resolve calcium signaling dynamics in freely behaving *Hydra*.

## 2 Methods

### 2.1 Microscope design

To image an entire *Hydra* at sufficient spatial resolution to resolve every neuron, we used a system like that described in Liu et al., 2022 [Figure 1]. Briefly, we used a 16× water immersion objective (Nikon, #MRP07220) with a tube lens of focal length *f*_*TL*_ = 300 mm and a Fourier lens of focal length *f*_*FL*_ = 200 mm, set up as a relay lens pair. We used a custom-machined 3×3 microlens array with a focal length *f*_MLA_ = 30 mm. 6 mm diameter MgF_2_-coated lenses (Edmund Optics #45-120) were cut into square lenses with 3.3 mm side length and mounted in a custom lens holder made from black acrylic. Images were acquired using an Andor Zyla 4.2. This system provides a 900 µm diameter field of view, with a theoretical lateral and axial resolution of approximately 3 µm and 6 µm, respectively (Liu et al., 2022)

**Figure 1:**
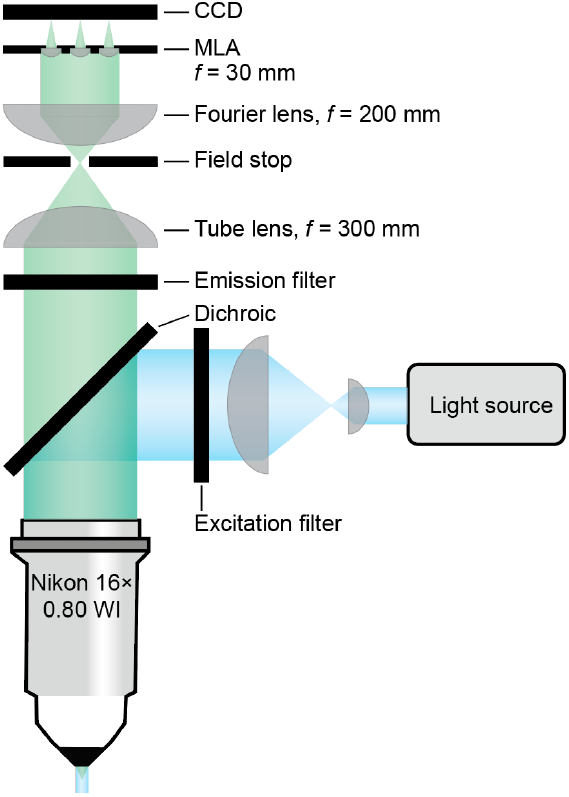
Microscope design. Schematic of the Fourier light field microscope.

### 2.2 Point spread function and calibration

Accurate knowledge of the point spread function is the most critical determinant of reconstruction quality in FLFM. Unlike conventional microscopy, where a poorly characterized PSF reduces resolution, errors in the FLFM PSF alter the entire reconstruction, misplacing signals axially, introducing ghost features, and amplifying noise. The PSF, therefore, must be measured carefully, with sufficient signal-to-noise to faithfully represent the optical response of each lenslet.

The PSF in light field microscopy is inherently three-dimensional and spatially variant. The shape of the PSF in each of the lenslets takes the form of an Airy disk in the focal plane, whose radius is set by the numerical aperture of the lenslet (Incardona et al., 2023). However, each of the lenslets subtends a different sub-aperture in the objective back focal plane, producing a distinct view of the Airy disk from a different angle. Consequently, the PSF of each lenslet is tilted with respect to the optical axis by an amount determined by the lenslet’s position in the array. In practice, a purely calculated PSF fails to capture aberrations, misalignments, and the precise tilt of each lenslet’s response. An experimentally measured PSF captures these effects, but raw measurements from a single bead suffer from a low signal-to-noise that may introduce noise-driven artifacts into the reconstruction.

We used a Hybrid PSF construction that combines experimental measurement with an analytical model (Liu et al., 2022). We imaged hundreds of beads (TetraSpeck Microspheres 1µm) embedded in 2% agarose by scanning the objective through the full axial range of the system (2 mm). This produces a z-stack in which many individual beads appear as diffraction-limited spots within each of the lenslet’s sub-images. These beads were then detected independently in each lenslet using a Laplacian-of-Gaussian (LoG) detector [Figure 2A]. The threshold for detection is automatically scaled by the intensity of each lenslet, to account for any systematic variation in lenslet illumination across the array.

**Figure 2:**
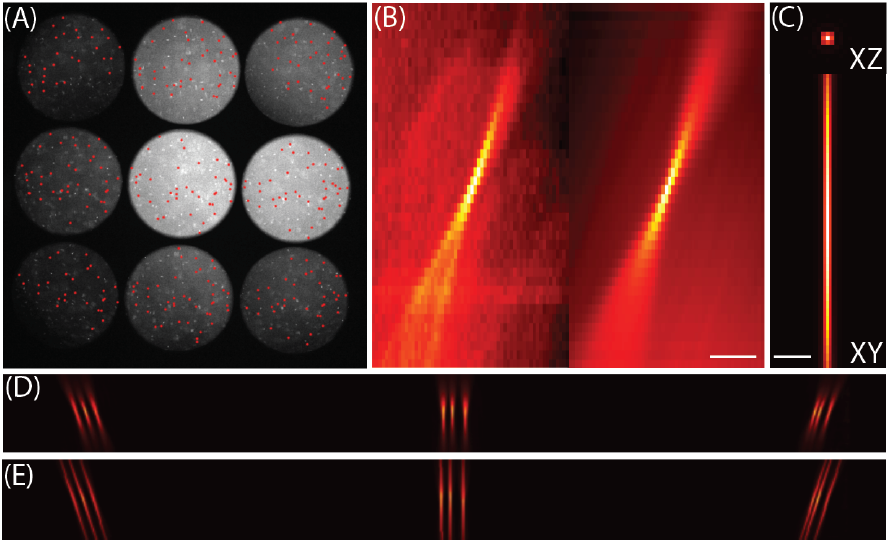
Making Hybrid PSF. (A) Detected beads overlaid on a single z-place of dense beads (center slice, z=145) of the bead stack; bead candidates identified by multi-scale blob detection (σ = 2.0–8.0 px, threshold = 0.01)). (B) Representative individual bead detections (left) and the corresponding depth-registered averaged PSF for one lenslet (right). (C) Simulated diffraction-limited PSF generated from an Airy-disk model of the optical system, shown as XZ (left) and XY maximum-intensity projection (right). (D) Full-field experimental PSF, assembled by placing each lenslet’s background-subtracted, intensity-weighted averaged bead PSF at its calibrated position and depth and normalizing to unit sum. (E) Final hybrid PSF formed by linearly blending the experimental (D) and simulated (C) PSFs (α = 0.15, weighting the experimental component), combining the empirically measured tilt and aberrations with the lower-noise diffraction-limited core. Scale bar: 25 µm

Detected beads were subjected to an isolation criterion, requiring a minimum separation from all neighboring detections by 50 µm in both the lateral and the axial dimensions, to ensure only unambiguously isolated beads contributed to the average. For each accepted bead, a three-dimensional patch of fixed lateral and axial extent was extracted (100 µm x 100 µm x 200 µm), centered on the bead position to sub-pixel accuracy using intensity-weighted centroid refinement to find the center point accurately [Figure 2B, S1]. The resulting average was computed independently for each of the nine lenslets, yielding nine individual lenslet PSF volumes with substantially improved signal-to-noise relative to an individual bead measurement [Figure 2B]. Beads were then matched between lenslets via cross correlation with the central lenslet [Figure S2], to determine the relative position between the nine PSF components to produce a single high-quality experimental PSF.

Rather than only using the averaged experimental PSF directly in the reconstruction, where residual noise can be amplified by iterative deconvolution, we used a computed Airy disk to generate a hybrid PSF. For each lenslet, a Gaussian fit was applied to the averaged PSF to determine the tilt axis of the lenslet’s response. A three-dimensional Airy disk was computed, using specified optical parameters of the system [Figure 2C]. This produces a smooth, noise-free PSF that is physically accurate in its lateral profile, correctly oriented in three dimensions. The nine per-lenslet PSFs are then assembled into a single full-field computed PSF [Figure 2D]. To allow flexibility between the experimental measurements and the analytical model, the final hybrid PSF was computed as the weighted combination

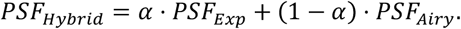

This parameterization allowed *α* to be adjusted continuously (Liu et al., 2022). We used *α* = 0.85 for reconstruction so that our PSF was primarily experimental [Figure 2E] but found little difference in reconstruction quality with varying *α* [Appendix 6.2].

### 2.3 Development of the reconstruction software

Volumetric reconstruction from an FLFM image is formulated as an inverse problem. A two-dimensional image *I(x, y)* formed on the detector is modeled as the sum of contributions from each axial plane of the sample volume, each convolved with the depth-dependent point spread function:

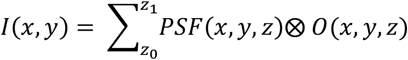

where *I* is the image on the CCD, PSF is the point spread function, *O* is the object in real space, and ⨂ denotes the two-dimensional convolution. Given the measured image *I(x, y)* and the PSF, the object *O*(*x, y, z)* can be recovered iteratively using a multiplicative Richardson-Lucy deconvolution:

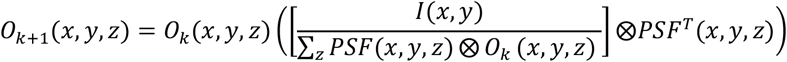

which is then iterated in *k* until sufficient convergence is reached, typically within 10-20 iterations for sparse samples. To characterize the convergence of our algorithm, we constructed a synthetic fluorescence volume containing 80 randomly placed 3D Gaussian blurs. We then simulated the corresponding raw FLFM sensor image by forward projecting the volume through the measured hybrid PSF; the exact operation the iterative reconstruction inverts. We ran the deconvolution for 40 iterations, tracking the relative update norm, 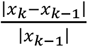 as the convergence criterion.

Convergence was defined as the first iteration at which this quantity fell to 5% of its value at iteration 1, yielding a threshold of approximately 15 iterations [Figure 3A], which produced volumes that were visually similar to the ground truth volume [Figure 3B, C]. All subsequent reconstructions were performed using 15 iterations.

**Figure 3:**
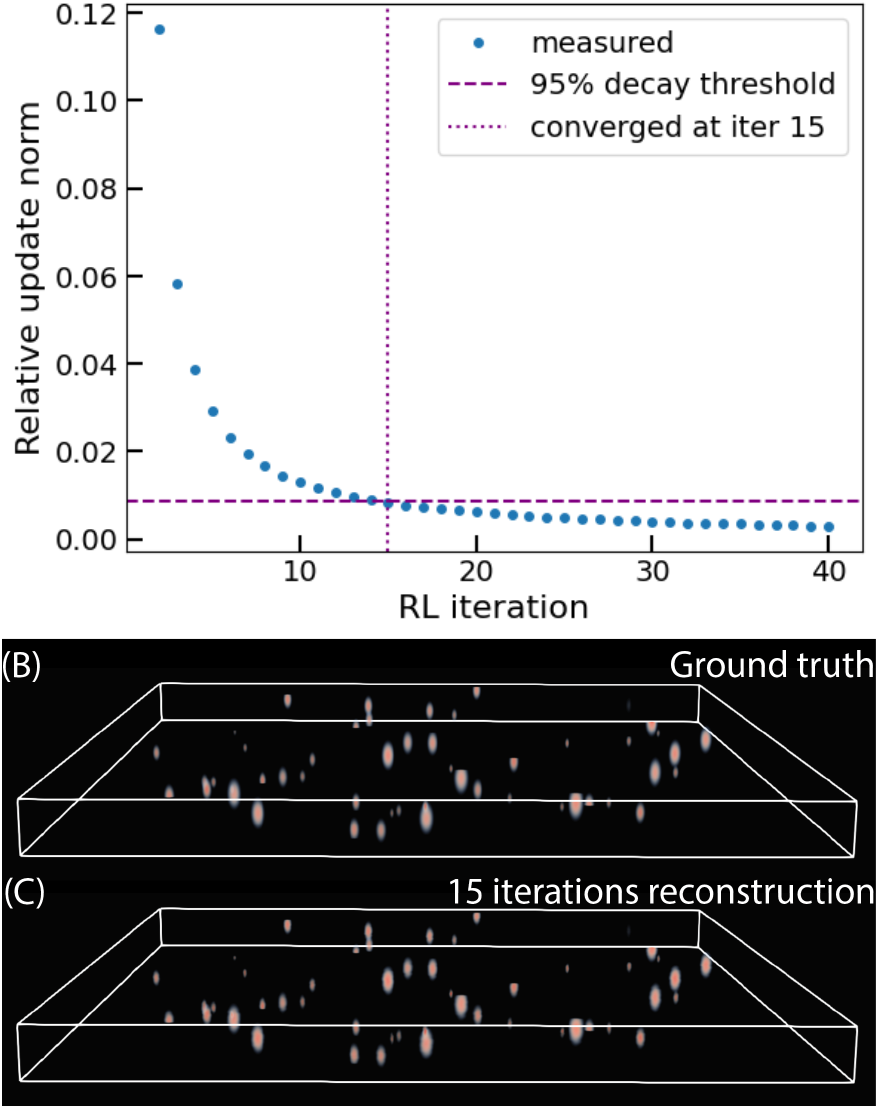
OpenFLR reconstructions. (A) Plot of the normalized average change in the deconvolved image at each iteration. By 15 iterations, the image has reached 95% of its convergence. (B) Generated ground truth image and (C) reconstruction after 15 iterations shows good agreement. Bounding box: 900 µm x 900 µm x 200 µm

The dominant computational cost in each iteration is the set of two-dimensional convolutions, which we computed in the Fourier domain. We performed several computational optimizations to improve the speed of the algorithm. A full description of the computational manipulations is given in Appendix 6.4. We tested three different implementations of the same underlying algorithm; numpy (no GPU acceleration), JAX and PyTorch. Additionally, we tested six different types of hardware: a MacBook Pro M2 CPU, and five different GPUs, the Quadro P5000, RTX A4000, V100, A100 and H100. All tests were done on a 2048 × 2048-pixel image with 40 depth planes at 15 iterations each. All convolutions were implemented using GPU-accelerated batch FFTs via PyTorch, with the PSF frequency-domain representation pre-computed and held resident on the GPU between frames.

First, we tested the PyTorch algorithm across all six architectures and found a significant increase in speed when moving from CPU to GPU (from 2-15 s / iteration to 89 ms/iteration). Higher-tier GPUs give us faster results [Table 1], with the best being an NVIDIA H100, with 6.7 ms per iteration. Next, we tried the same algorithm, implemented in Python in several ways. We found a range of reconstruction times, with the slowest being pure numpy and the fastest being JAX at 6.3 ms per iteration [Table 1]. The latter yields a mean reconstruction time of approximately 95 ms per image (15 iterations), corresponding to an average frame rate of approximately 10.5 fps. This exceeds the temporal requirements imposed by GCaMP6s calcium dynamics in *Hydra*, which evolve on timescales of approximately 500 ms, and is sufficient for real-time display and processing during acquisition.

**Table 1.** Speed of reconstruction. Time for a single iteration of reconstruction, taken as the average of 20 iterations. Benchmarking the same algorithm against three software and six hardware implementations.

|  | MB M2 | Quadro P5000 | RTX A4000 | V100 | A100 | H100 |
| --- | --- | --- | --- | --- | --- | --- |
| JAX |  |  |  | 23.6 ms | 11 ms | 6.3 ms |
| PyTorch | 2.0 s | 89 ms | 48 ms | 22.4 ms | 11 ms | 6.7 ms |
| numpy |  |  |  | 4.9 s | 15.6 s | 2.8 s |

While JAX was the fastest reconstruction implementation, it is limited to running on GPUs with Unix systems. PyTorch, on the other hand, can run on any system and is able to fall back to CPU when a GPU is not found. This flexibility makes it an attractive alternative. PyTorch yielded a mean reconstruction time of approximately 100 ms per image, corresponding to an average frame rate of approximately 10 fps, still well within the limit needed to be truly ‘real-time’ for our use case.

### 2.4 Three-dimensional particle tracking via multi-view triangulation

For sufficiently sparse samples, full volumetric reconstruction is not always necessary to recover particle trajectories. We therefore developed a reconstruction-free localization approach that estimates three-dimensional particle positions directly from raw light field frames by triangulating each emitter across the lenslet sub-apertures, enabling tracking at frame rates substantially higher than is achievable by reconstruction.

This approach exploits the parallax intrinsic to the light field. A fluorescent bead at position (*x*_0_, *y*_0_, *z*_0_) appears in each of the lenslet sub-apertures as a spot whose lateral position depends on both the emitter’s planar position and its depth. Because each sub-aperture views the sample from a different angle, the spot in the i-th lenslet is displaced from the emitter’s nominal lateral position by an amount that varies linearly with depth, *p*_*i*_(*x*_0_, *y*_0_, *z*_0_) = (*x*_0_, *y*_0_) + ***c***_***i***_ + ***m***_***i***_(*z* – *z*_0_), where ***c***_***i***_ is the offset between lenslets and ***m***_***i***_ = (*m*_*x*,,*i*+_, *m*_*y,i*_) is the per-lenslet parallax slope. The slopes ***m***_***i***_ and offset can be measured by the calibration: the hybrid PSF construction already fits the axial trajectory of each lenslet’s spot, we read the parallax slope from the same fit used to build the reconstruction PSF, ensuring the localization and reconstruction share a single consistent geometric calibration.

Given these slopes, the depth of an emitter is the only remaining unknown once its lateral position is fixed, and a hypothesized position (*x, y, z)* predicts the location in every sub-aperture. We used this constraint to recover emitters without ever forming a volume. Candidate spots were first detected in each sub-aperture by matched filtering the background subtracted image against the calibrated PSF and extracting local maxima, refined to sub-pixel accuracy by parabolic interpolation. Correspondence and depth were then solved jointly. For each candidate spot in the central sub-aperture, we swept the depth z over the calibrated range. At each depth, the parallax model predicts the spot position in every other sub-aperture; we counted those in which a detected spot fell within a fixed tolerance. The depth maximizing this agreement identifies both the emitter’s axial position and the set of sub-apertures in which it is observed. An emitter was accepted when it was triangulated by a minimum of five sub-apertures, allowing its position to be refined by solving the overdetermined system of 2N parallax equations for N views, in a single least-squares step. This consensus-based correspondence search follows the logic of random sample consensus like RANSAC (Fischler and Bolles, 1981), with the minimal hypothesis enumerated over the calibrated depth range rather than sampled at random. Because the geometry constrains which spots can correspond, this search scales with the number of spot pairings, and near-coincident solutions were merged so that each emitter is reported once.

The multi-view consensus makes this method more robust than matched filtering alone. An emitter produces a consistent parallax track across many sub-apertures, whereas a spurious detection of an accidental pairing satisfies the pairing in only a few. Requiring agreement across a majority of views rejects false positives that no single sub-aperture could exclude. The same consensus resolves moderate crowding: two emitters that overlap in one sub-aperture generally separate in others due to their differing depths which carry their images along different parallax tracks.

Per-frame coordinates were then linked into continuous three-dimensional trajectories using the Crocker–Grier nearest-neighbor algorithm as implemented in TrackPy (Allan et al., 2025; Crocker and Grier, 1996), with a maximum inter-frame displacement of 10 pixels and a gap-closing memory of 5 frames. All tracks were required to persist for 20 frames to be considered real tracks. This produced clear tracks of the point-like particles [Figure 4A]. To validate the localization, we applied the method to synthetic light field frames generated by convolving beads at known three-dimensional positions with the measured PSF, so that the recovered positions could be compared against ground truth (Appendix 6.3). Because the frames are formed by PSF convolution while positions are recovered by parallax triangulation, this test is independent of the model being validated and confirms that the method recovers the true positions of well-separated emitters to sub-voxel accuracy at densities comparable to our experimental recordings.

**Figure 4:**
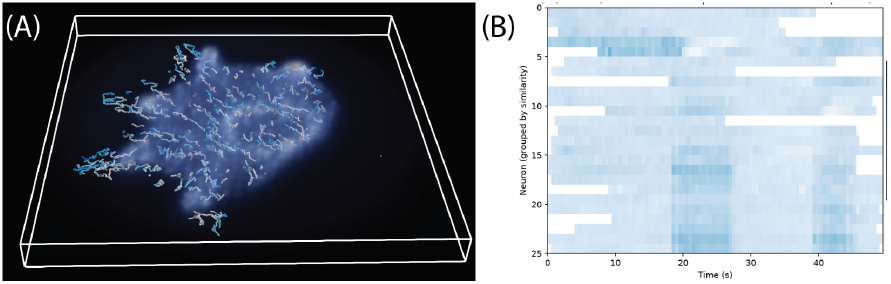
Tracking neurons using Fourier light microscopy. (A) Linked neuron trajectories overlaid on the reconstructed volume of the tdTomato channel, demonstrating that identity is maintained across large three-dimensional body deformations. (B) GCaMP ΔF/F heatmap, with neurons sorted by the time of their peak response. Bounding box: 900 µm x 900 µm x 200 µm

With the matched-filter kernel pre-computed and resident on the GPU and the sub-aperture convolutions evaluated by batched two-dimensional FFTs, detection and triangulation required 75 ms per frame on a low-tier GPU (Quadro p5000) and 214 ms on a CPU, roughly an order of magnitude faster than a full reconstruction would take. Because the emitter’s depth is recovered from parallax across many sub-apertures rather than from a single reconstructed volume, localization remains reliable even though no iterative deconvolution is performed, at a small fraction of the cost of the full reconstruction.

### 2.5 *Hydra* as a model system for testing OpenFLR

To test the utility of our software, we used the light-field microscope to image and reconstruct the activity of the nervous system of *Hydra vulgaris. Hydra* is an emerging system in neuroscience, due to its sparse nervous system spread out over its body, enabling simultaneous imaging of its entire nervous system (Dupre and Yuste, 2017). We chose small *Hydra* with approximately 300 neurons and an 800 µm long body. We used transgenic *Hydra* expressing both tdTomatoNLS and GCaMP6s in their interstitial cell layer (Hanson et al., 2023). Imaging two channels lets us track each neuron on the constitutive tdTomato signal, which is present independent of activity, while reading calcium dynamics from the GCaMP channel, so that transient loss of activity does not break a trajectory. To accurately track the activity of every neuron, we needed to image each one at a time resolution on the order of hundreds of milliseconds. This offers an ideal test case for FLFM and the OpenFLR reconstruction software; a sparse, large, and rapidly evolving system of point-like particles.

To recover neural activity, we localized and tracked neurons on the structural tdTomato channel and then read out calcium dynamics from the GCaMP channel at the tracked positions. For each frame, three-dimensional peaks were detected in the matched-filter score volume with a Laplacian-of-Gaussian blob detector and linked across frames using TrackPy (search range 10 px, memory 5 frames), retaining trajectories spanning at least 20 frames. Because tdTomato is expressed constitutively, these trajectories persist even when a neuron is transiently silent, so each neuron’s identity is maintained throughout the recording regardless of its activity. At every tracked position, we then registered the coordinate into the GCaMP channel and sampled the fluorescence within a small circular ROI (4-px radius), and computed the relative change in fluorescence *ΔF/F* = (*F − F*_0_)/*F*_0_, taking the 20th percentile of each trace as the baseline F_0_. The resulting traces resolved clear calcium transients across the 112 tracked neurons, several of which were co-active within single frames [Figure 4B], revealing synchronous firing across the nerve net that would be inaccessible to a single-plane or reconstruction-only workflow.

### 2.6 Software architecture and availability

OpenFLR is intended to be modular and open source. The code is implemented in Python and organized around a single utility module containing the calibration, reconstruction, localization, and tracking routines, together with a configuration module that specifies the optical parameters of the system. All compute-intensive operations, namely the Richardson-Lucy forward and transpose convolutions and the matched-filter cross-correlations, are expressed as FFTs in PyTorch, so the same code runs on either CPU or GPU without modification and automatically uses CUDA when a compatible device is available. Particle linking is delegated to TrackPy.

The reconstruction pipeline has four interchangeable front ends that share this common core: an annotated Jupyter notebook that walks through the full workflow from calibration to trace extraction; a command-line script for unattended batch processing; an interactive web application for users who prefer a graphical workflow with live parameter tuning; and an ImageJ/Fiji plugin that brings reconstruction into a tool already familiar to most microscopists. Because the optical configuration is fully parameterized, the same code supports microlens arrays of arbitrary size and pitch rather than being hard coded to the 3 × 3 array used here. All optical parameters are defined in a separate Python file and can be modified by the user prior to running the code.

The FLFM tracking routine and calcium signal extraction are available in the Jupyter Notebook, with interactive plots for tuning the detection threshold and peak distance cutoff and for inspecting bead detection. Because reliable tracking depends on a small number of sample-specific choices that are best made by eye, this human step is most effective in the notebook environment, rather than an unattended or graphical workflow. All four interfaces, together with example data and the calibration routines, are available at GitHub.com/OpenFLR, and data sufficient to reproduce the results in this work are deposited on Dryad [Adkins et al. 2026].

## 3 Discussion

Here, we have shown the utility of the OpenFLR software package for advancing Fourier lightfield microscopy. This software was able to reconstruct volumes effectively in real time. OpenFLR is easily adaptable for lens arrays of variable sizes and lenslet number, without affecting the reconstruction speed per iteration. In addition, it can serve as a basis for other orthographic-based imaging modalities, like XLFM or Fourier DiffuserScope (Cong et al., 2017, Liu et al., 2020).

Several limitations bound the present approach. The matched-filter localization assumes that each emitter is well approximated by a shifted copy of the calibrated PSF; it is therefore best suited to point-like sources, and we expect its performance to degrade as emitter density increases and points begin to overlap, at which point full reconstruction remains the more appropriate route. Like all FLFM systems, axial resolution is poorer than lateral resolution, and both depend on the accuracy of the PSF calibration. Reconstruction quality also relies on the assumption that the PSF is laterally invariant within each sub-aperture, which holds only over a limited field of view. Finally, the speed figures reported here are specific to our hardware and operating parameters. Performance on other systems will scale with available GPU memory and the chosen volume size, depth range, and iteration count.

## Supporting information

Supplemental Infromation

