## Supplemental Infromation for "Rapid volumetric reconstruction and tracking for Fourier light-field microscopy enables real-time calcium imaging in freely behaving *Hydra*"

1.1 *Hydra* culture

We cultured adult *Hydra* according to previously established protocols (Dupre and Yuste, 2017). Briefly, *Hydra* vulgaris (AEP) were cultured at 18°C in standard *Hydra* media (1.3 mM CaCl2, 0.2 mM MgCl2, 0.02 mM KNO3, 0.5 mM NaHCO3 and 0.8 mM MgSO4). *Hydra* were kept on a 12h: 12h light: dark cycle. Adult polyps were fed three times each week with freshly hatched brine shrimp (Artemia Nauplii, Great Salt Lake).

1.2 Sensitivity of reconstruction to $\boldsymbol{\alpha}$

To determine the sensitivity of the reconstruction to our choice of α, we reconstructed a single light-field frame using hybrid PSFs spanning the full range of the blending parameter (α = 0, 0.2, 0.4, 0.6, 0.8, and 1.0) and quantified how much each reconstruction differed from a reference volume of α = 0.85. Because Richardson–Lucy reconstructions can differ in absolute intensity scale, we compared volumes using the Pearson correlation coefficient, which is invariant to affine rescaling of voxel intensities and therefore reports structural agreement independent of overall brightness. Correlations were computed voxel-wise over the cropped image region [Table S1]. Across the entire range of α, reconstructions remained highly correlated with the α = 0.85 result (Pearson r ≥ 0.992), with the largest deviation occurring for the pure-simulated PSF (α = 0, r = 0.992).

| Reconstruction (vs. α = 0.85, 15 iter) | Pearson r |
| --- | --- |
| α = 0 | 0.993 |
| α = 0.2 | 0.995 |
| α = 0.4 | 0.997 |
| α = 0.6 | 0.999 |
| α = 0.8 | 1.000 |
| α = 1.0 | 0.992 |
| α = 0.85, 10 iterations | 0.993 |
| α = 0.85, 20 iterations | 0.993 |
| α = 0.85, 40 s later time point | 0.656 |

To place this variation on a meaningful scale, we compared it against two references. First, we reconstructed the same frame at 10 and 20 RL iterations (versus the default of 15) and correlated these with the α = 0.85, 15-iteration reference; both yielded r ≈ 0.993, comparable to the variation introduced across the full α range. Because 15 iterations already brings the reconstruction to within 5% of convergence [Figure 3A], the effect of α is no larger than the residual variation associated with the algorithm's own convergence tolerance. Second, to confirm that the Pearson coefficient is in fact sensitive to genuine structural differences between these volumes, we correlated our reference volume with a later time point (40 s apart), which gave r ≈ 0.65.

Taken together, these results show that the precise value of the blending parameter has a negligible effect on the reconstruction: the variation across the entire range of α is comparable to the algorithm's convergence tolerance and significantly smaller than the frame-to-frame variation of the sample itself. We therefore fixed α = 0.85 for all reconstructions, but the pipeline is robust against this choice.

1.3 Validation of triangulation localization against ground truth

Because the true positions of emitters in experimental recordings are unknown, we validated the localization on synthetic data for which the ground truth is defined. Synthetic light field frames were generated by placing beads at known positions and forming each frame through the same image-formation model as the real instrument: convolving the synthetic volume at each depth plane with the point spread function and flattening the image. The images are then corrupted with shot noise, motion blur, and background haze.

At an emitter density comparable to our experimental recordings (75 beads across the 220 x 220 x 36 voxel volume), the method recovered well-separated emitters with a localization precision of 82% and a per-frame detection recall of 78%, with the estimated emitter count matching the true count. Localization error was strongly bimodal: the majority of beads were recovered to within a fraction of a voxel of their true positions while a minority, like those overlapping other emitters closely enough that their sub-aperture spots could not be separated, were localized incorrectly. This behavior is consistent with the point-source assumption underlying the method: an emitter is recovered accurately when its parallax track can be distinguished from those of its neighbors and fails when it cannot. As emitter density increases (150 beads across the 220 x 220 x 36 voxel volume), the fraction of beads whose parallax tracks overlap grows, and recall declines accordingly; at the higher densities, characteristic of dense labeling, a substantial fraction of emitters cannot be separated, and full volumetric reconstruction becomes the more appropriate route. All simulated images are available on Dyrad [Adkins et al. 2026].

1.4 Reconstruction optimization

Starting from the general RL deconvolution equation,

$$O_{k+1}(x,y,z)=O_{k}(x,y,z)\left( \left[ \frac{I(x,y)}{\sum_{z} PSF(x,y,z)\bigotimes O_{k}(x,y,z)} \right]\bigotimes PSF^{T}(x,y,z) \right)$$

We use the convolution theorem to write

$$\left[ \frac{I(x,y)}{\sum_{z} PSF(x,y,z)\bigotimes O_{k}(x,y,z)} \right]\bigotimes PSF^{T}\left( x,y,z \right)=\mathcal{F}^{-1}\left\{ \mathcal{F}\left\{ \frac{I\left( x,y \right)}{\sum_{z} \mathcal{F}^{-1}\mathcal{\{F}\left\{ PSF\left( x,y,z \right) \right\}\mathcal{\cdot F\{}O_{k}\left( x,y,z \right)\}\}} \right\}\mathcal{\cdot F\{}PSF^{T}\left( x,y,z \right)\} \right\}$$

We can then move the sum inside the inverse Fourier transform

$$\mathcal{F}^{-1}\left\{ \mathcal{F}\left\{ \frac{I\left( x,y \right)}{\mathcal{F}^{-1}\{ \sum_{z} \left( \mathcal{F}\left\{ PSF\left( x,y,z \right) \right\}\mathcal{\cdot F}\left\{ O_{k}\left( x,y,z \right) \right\} \right) \}} \right\}\mathcal{\cdot F\{}PSF^{T}\left( x,y,z \right)\} \right\}$$

which eliminates all but one inverse Fourier transforms in the denominator. We can then compute the Fourier transform of the PSF and the transpose of the PSF outside of the function, defining these to be $PSF_{FFT}\mathcal{= F}\left\{ PSF\left( x,y,z \right) \right\}$, $PSF_{FFT}^{T}\mathcal{= F}\left\{ PSF^{T}\left( x,y,z \right) \right\}$. The equation then becomes

$$\mathcal{F}^{-1}\left\{ \mathcal{F}\left\{ \frac{I\left( x,y \right)}{\mathcal{F}^{-1}\{ \sum_{z} \left( PSF_{FFT}\mathcal{\cdot F}\left\{ O_{k}\left( x,y,z \right) \right\} \right) \}} \right\}\cdot PSF_{FFT}^{T} \right\}$$

So, then each iteration becomes

$$O_{k+1}\left( x,y,z \right)=O_{k}\left( x,y,z \right) \mathcal{F}^{-1}\left\{ \mathcal{F}\left\{ \frac{I\left( x,y \right)}{\mathcal{F}^{-1}\{ \sum_{z} \left( PSF_{FFT}\mathcal{\cdot F}\left\{ O_{k}\left( x,y,z \right) \right\} \right) \}} \right\}\cdot PSF_{FFT}^{T} \right\}$$

Which means that, in each reconstruction step, we only require one 3D FFT, two matrix element multiplication steps, one matrix element division, one 2D FFT and one 2D iFFT.

1.5 Code and data availability

To ensure this code could be widely utilized as much as possible, we made the code available as a Python notebook, Python script, dash application, and as an ImageJ plugin. Each of these is available at GitHub.com/OpenFLR. Installation instruction and version information is available via the GitHub. Data sufficient to reproduce all results from this paper are hosted on Dryad [Adkins et al. 2026].

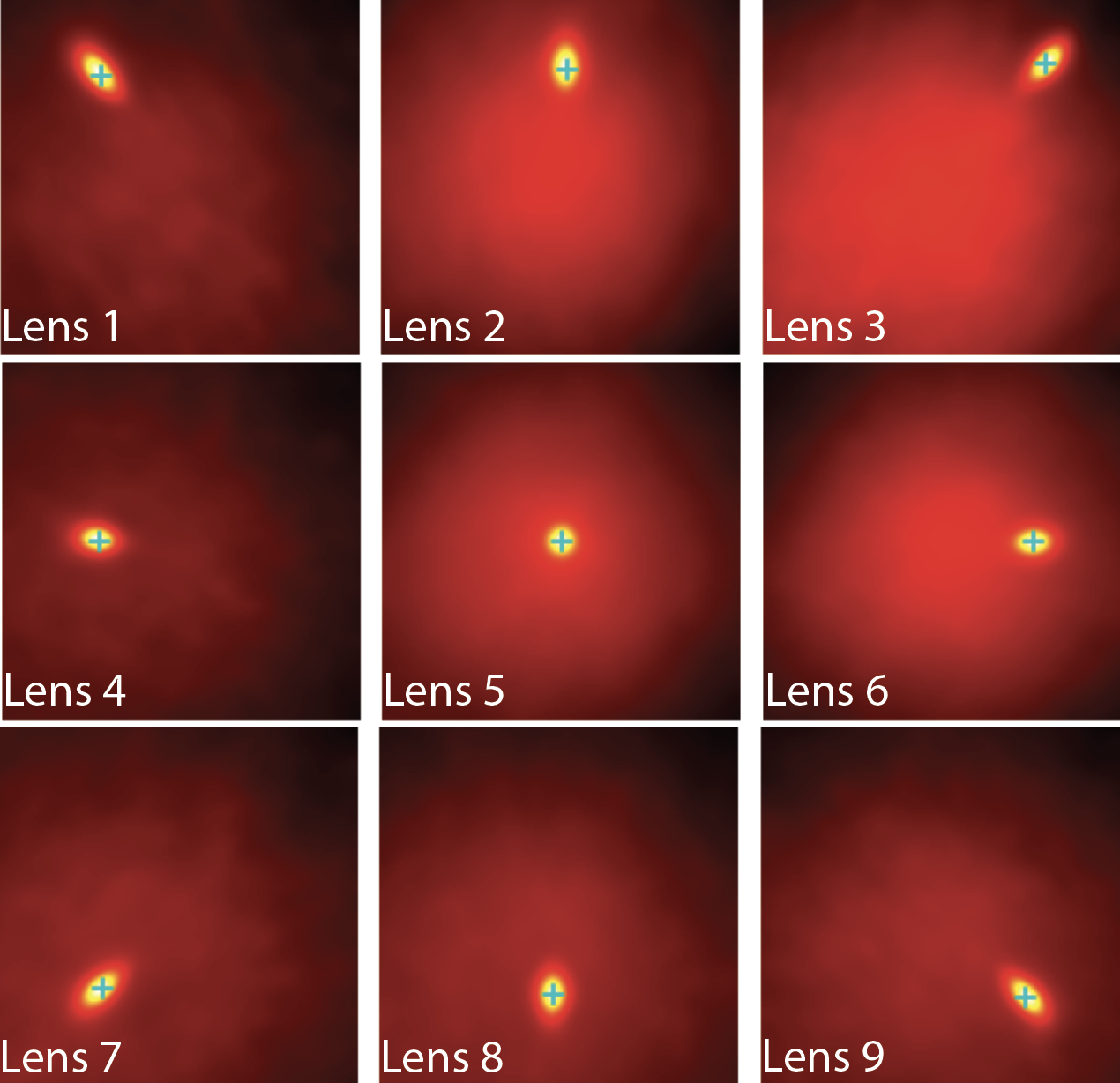

**Figure S2:** Cross-correlation between the averaged bead PSF of each lenslet and the center lenslet, computed over a ±60 px search window (signal threshold = 0.01), used to determine the relative spatial offset between lenslets prior to PSF assembly.

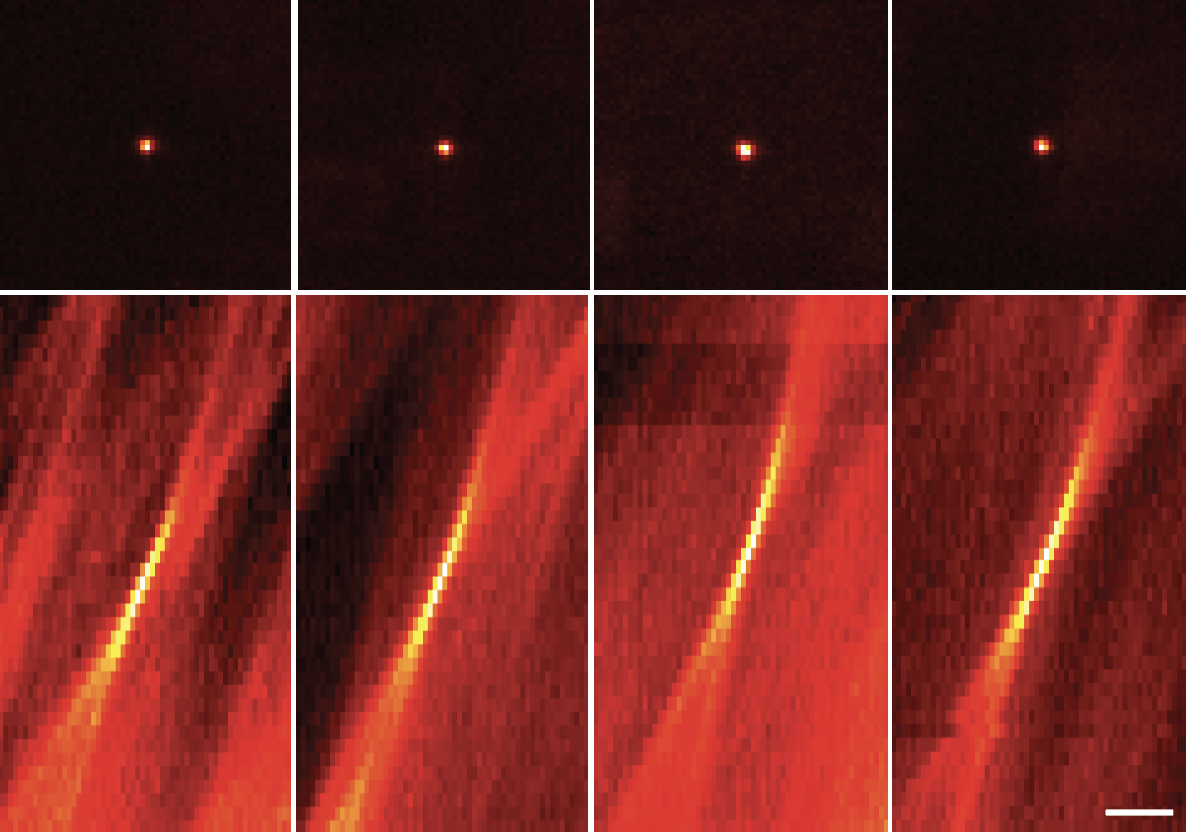

**Figure S1:** Four example beads detected within lenslet 4, shown as an XY image through the bead centroid (top row) and an XZ projection (bottom row). Beads were identified by blob detection (σ = 2.0–8.0 px, threshold = 0.01) within a 3D bounding box (100 x 100 x 200), were required to be and aligned to a common center before averaging. Scale bar: 25 µm
